# Mapping cell types in the tumor microenvironment from tissue images via deep learning trained by spatial transcriptomics of lung adenocarcinoma

**DOI:** 10.1101/2023.03.04.531083

**Authors:** Kwon Joong Na, Jaemoon Koh, Hongyoon Choi, Young Tae Kim

## Abstract

Profiling heterogeneous cell types in the tumor microenvironment (TME) is important for cancer immunotherapy. Here, we propose a method and validate in independent samples for mapping cell types in the TME from only hematoxylin and eosin (H&E)-stained tumor tissue images using spatial transcriptomic data of lung adenocarcinoma. We obtained spatial transcriptomic data of lung adenocarcinoma from 22 samples. The cell types of each spot were estimated using cell type inference based on a domain adaptation algorithm with single-cell RNA-sequencing data. They were used to train a convolutional neural network with a corresponding H&E image patch as an input. Consequently, the five predicted cell types estimated from the H&E images were significantly correlated with those derived from the RNA-sequencing data. We validated our model using immunohistochemical staining results with marker proteins from independent lung adenocarcinoma samples. Our resource of spatial transcriptomics of lung adenocarcinoma and proposed method with independent validation can provide an annotation-free and precise profiling method of tumor microenvironment using H&E images.

## INTRODUCTION

The tumor microenvironment (TME), which comprises stromal and immune cells, is associated with tumor initiation, progression, and metastasis ^1-4^. Notably, the importance of TME in cancer treatment has been strengthened since immune checkpoint inhibitors (ICIs) have dramatically changed cancer treatment ^5, 6^. However, only a subset of patients responds to ICIs, while the others eventually experience disease progression. One of the reasons for the resistance or non-response to ICI is the immunosuppressive TME ^7, 8^. Therefore, various studies have been conducted to decipher the intricate components of the TME that impact response to ICIs. These studies have aimed to improve the accuracy of response prediction and to design new treatments that various targets such as stromal and immune cells within the TME overcoming the limitations associated with ICI resistance ^9, 10^.

The spatial patterns of various cell types in the TME are key factors in predicting therapeutic response and prognosis ^11-13^. Single-cell RNA sequencing (scRNA-seq) approaches provide molecular information to decipher cell types in the TME, but one of the drawbacks is the loss of spatial information. Spatial transcriptomics is a new technology that obtains whole-genome expression data from numerous small spots within tissue ^14^. Since this method produces histopathological images as well as gene expression data, it can provide an opportunity for an integrative analysis strategy of imaging and gene expression to understand the spatial patterns of TME ^15, 16^. Besides, analysis of the whole-slide histologic images with deep learning (DL) algorithms has become a significant and rapidly growing tool in analyzing the spatial heterogeneity of TME ^17^. This approach could also provide predicted spatial information on cell types, which include immune cells, molecular markers such as programmed death-ligand 1 (PD-L1), and cancer cells ^18-20^. Considering multiple cell types could be defined by spatial transcriptomics with high resolution, training DL on tissue images with these multiple cell types could replace manual labeling by experts that need intensive and time-consuming process ^17^.

In this study, we present a strategy of cell-type mapping to understand spatial patterns by using spatial transcriptomic data and hematoxylin and eosin (H&E) histopathological images. We obtained spatial transcriptomic data with H&E images of lung adenocarcinoma (LUAD) to develop the model, which captured the spatial distribution of multiple cell types in the TME. This model was validated in a separate set of spatial transcriptomic data. We also found that the DL model effectively predicted the spatial distribution and proportion of cell types in independent LUAD samples using an immunohistochemistry (IHC) assay as an external validation study. With this study, we provide a resource of spatial transcriptomic data of LUAD and present a framework for producing DL models to effectively map cell types from H&E images without the need for human labeling for H&E image-based biomarkers.

## METHODS

### Patients Samples

Overall, 22 LUAD samples were collected from lung specimens of 10 patients who underwent curative surgical resection. The collected samples were embedded in an optimal cutting temperature compound (25608-930, VWR, USA) in the operating room and stored at -80°C until cryosectioning. For the cryosectioning, samples were equilibrated to -20°C with a cryotome (Thermo Scientific, USA) and cut to a thickness of 10 μm. Next, the sections were imaged and processed for spatially resolved gene expression using the Visium Spatial Transcriptomic kit (10X Genomics, USA). The study protocol was reviewed and approved by the Institutional Review Board of Seoul National University (application number: H-2009-081-1158). **Supplemental Table S1** summarizes the clinical information of all the patients.

### Slide preparation and RNA sequencing

First, tissue sections were placed on chilled Visium tissue optimization slides (1000193, 10X Genomics, USA) and Visium spatial gene expression slides (1000184, 10X Genomics). Next, the sections were fixed in chilled methanol and stained with H&E, and the brightfield images of the H&E slides were obtained. For the gene expression analysis, cDNA libraries were constructed following the Visium Spatial Gene Expression User Guide. Briefly, the sections were incubated with permeabilization enzymes, and the timing was determined using the Visium Spatial Optimization Slides (1000193, 10X Genomics, USA). Subsequently, after washing with saline sodium citrate buffer, RT Master Mix (Visium Reagent kit, 10X Genomics, USA), which contained reverse transcription reagents, was added to the permeabilized tissue sections and reverse transcription was performed based on the manufacturer’s protocol. Next, after reverse transcription, the sections were incubated in KOH (0.08 M) for 5 min and then incubated in Second Strand Mix for 15 min at 65°C. Finally, cDNA amplification and library construction were performed. Sequencing was performed on a NovaSeq 6000 System S1 200 (Illumina, USA) at a sequencing depth of approximately 250 megadread pairs per sample.

### Processing of the raw spatial transcriptome data

Raw FASTQ files and histology images were processed using Space Ranger v1.1.0 software. The alignment process was based on the Spliced Transcripts Alignment to a Reference (STAR) v.2.5.1b, using the hg38 reference genome. Next, the alignment and count process were performed using the ‘spaceranger count’ command by specifying the input of FASTQ files, reference, section image, and Visium slide information. The pipeline detected the tissue area by aligning the image to the printed fiducial spot pattern of the Visium slide and recognizing stained spots from the image.

### Cell type score estimation from spatial transcriptomic data

Spatial transcriptomic data based on barcodes were used to estimate cell-type scores. Therefore, to define the cell types, scRNA-seq data obtained from human LUAD were used ^21^. Next, the cell types defined by the scRNA-seq data were transferred to the spatial transcriptomic data of the multiple LUAD samples using CellDART ^22^. We used 42 cell types of scRNA-seq data defined in a previous study (**Supplemental Table S2**). The markers for each cell type were selected using the Wilcoxon test in the Scanpy package (version 1.5.1) ^23^. In addition, for the CellDART, we selected 10 marker genes per cell type, and 10,000 pseudospots were generated with the number of mixed cell types set to 8. The cell type scores of each spot were estimated, and the five major cell type scores were calculated by the sum of subtype scores.

### A convolutional neural network model for predicting cell type scores

We utilized spatially registered spatial transcriptomic data co-registered with H&E images to develop a model that predicts cell type scores. Image patches were acquired using H&E slide images from the spatial transcriptomics after the stain normalization process using the staintools python package (https://pypi.org/project/staintools). The center of each image is similar to a spot, which represents the gene expression data. In addition, the sizes of the image patches were adjusted to have a 128 × 128 × 3 matrix size of approximately 4 μm per pixel. Therefore, image augmentation was included to train the model by using rotation, zooming with 20% range, RGB channel shift with 20% range, and random horizontal and vertical flips. Overall, 43,074 patches and corresponding spots obtained from the 19 samples were used for the model training. Among these patches, 5% of randomly selected patches were used for the internal model validation. Finally, 5,653 patches obtained from three sample were used as the independent dataset.

A convolutional neural network (CNN) model based on the ResNet50 backbone was constructed to estimate the cell type scores derived from the CellDART. The final layer of the ResNet50 model was excluded, and a global average pooling layer was added, followed by a fully-connected layer with 1024 dimensions. The final output layer had five nodes representing the five major cell type scores. An Adam optimizer with a learning rate of 1 × 10^−5^ was used for the optimization. Furthermore, the Poisson loss function was used as the loss function, which is defined as follows:

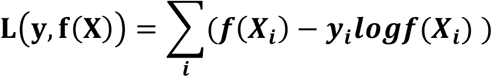

### DL-based cell types enrichment scores from H&E images

The model trained by the H&E images paired with spatial transcriptomic data was applied to the H&E images. For the application, the pixel sizes were adjusted to 4 μm, and stain normalization was conducted as the training set. In addition, the DL model was directly applied to estimate the score of the cell types for each patch in the datasets with H&E image patches. For the whole slide image, the trained DL models were repeatedly applied to the patches defined by the sliding windows to estimate the scores for the center of each window. Furthermore, for windowing to obtain image patches, the sliding step was set to 64 pixels for the whole slide images. Finally, each window size was set to 128 × 128.

### Correlation with cell type enrichment scores estimated by bulk RNA-seq of TCGA data

TCGA data comprised histopathological images and omics data across multiple cancer types. Histopathologic image data of 479 patients with LUAD who had also bulk RNA-seq were downloaded from GDC portal (https://portal.gdc.cancer.gov/). Next, 5 cell types enrichment score maps were estimated by the trained DL model and H&E images of TCGA. The 5 cell type scores of each sample were evaluated by mean value of whole tissue region. To compare the results of DL-based cell type scores with those of other methods, cell type enrichment scores were estimated from RNA-sequencing data. The xCell tool (http://xcell.ucsf.edu/) was used to infer cell types from tissue transcriptome profiles ^24^. In addition, to compare the T/NK cell enrichment score estimated by DL model, cytotoxic score was estimated ^25^. We estimated cell enrichment scores for B cells, macrophages and microenvironment score of xCell. Subsequently, we performed Spearman’s correlation analysis between cell type enrichment scores estimated from RNA-sequencing data and those from the histopathological images using our DL-based prediction model.

### H&E and IHC assay for external validation

Representative H&E-stained slides from 11 patients with LUAD who underwent curative surgical resection at our institution. The H&E-stained slides were scanned using a WSI scanner Aperio GT 450 WSI scanner (Leica Biosystems, Germany) at 40x the original magnification. **Supplemental Table S3** summarizes the clinicopathological characteristics of the patients. We predicted the cell enrichment scores of the five cell types from the H&E-stained slides using our model. The non-stained slices adjacent to the H&E-stained slides were taken. In addition, IHC staining for the following six markers was performed: CD3 (clone 2GV6, Ventana Medical Systems, Tucson, AZ, USA), CD20 (clone L26, DAKO, Carpinteria, CA, USA), CD56 (clone 123C3. D5, Cell Marque, Rocklin, CA, USA), CD68 (clone KP1, DAKO), pan-cytokeratin (clone AE1/AE3, DAKO), and SMA (clone 1A4, DAKO). Staining was performed using a Ventana Benchmark XT (Ventana Medical Systems, Inc., Tucson, AZ, USA) or Bond-Max autostainer (Leica Microsystems, Melbourne, Australia).

### External validation with surgically resected LUAD samples

Since H&E and IHC slides were obtained from the adjacent tissue sections, they were not completely registered with each other. For each IHC image, a deconvolution for DAB staining results was performed for all slides using the ‘rgb2hed’ function in the scikit-image package. Next, the spatial registration was performed using the symmetric diffeomorphic image registration ^26^. This process was performed using the DiPy Python package. In addition, both H&E and IHC slide images were converted to a grey scale. The registration process included linear registration followed by nonlinear registration. In addition, the center of mass was matched first, and the rigid and affine transformations were performed based on the mutual information of the two grayscale images. The nonlinear transformation was performed using a function, Symmetric Diffeomorphic Registration, with a cross-correlation similarity metric. The final warping matrix was estimated and applied to the deconvoluted image of the DAB channel. Therefore, the warped DAB channel image was co-registered with the corresponding H&E image. In contrast, the five cell-type maps predicted by H&E images were co-registered with the corresponding DAB channel image.

The cell type maps and the deconvoluted images for the DAB channel were subjected to Gaussian smoothing using a specific parameter, sigma = 10. Finally, a pixel-wise Spearman’s correlation analysis was performed to measure the similarity between DAB staining results and cell type maps predicted using the H&E images.

### Statistical analysis

Statistical analyses were performed using the R software package version 4.1.1 and Python with the SciPy package (version 1.5.1). The correlation between variables was evaluated using Pearson’s or Spearman’s correlation analysis, as applicable.

### RESULTS

### Model development and validation

**Figure 1** shows a schematic of the workflow of the proposed model. The H&E histopathological images were converted to patches corresponding to spots, which were classified into training and internal validation sets. We designed the model to predict five cell types (B cells, T/NK cells, myeloid cells, fibroblasts, and epithelial cells) of each spot in the H&E-stained histopathological image patch. First, we mapped five cell types using spatial transcriptomic data with cell types defined by the scRNA-seq of human LUAD ^21^. The cell type scores for each spot was estimated using a domain adaptation (CellDART) ^22^. The five major cell type enrichment scores were calculated as the sum of specific subtypes defined by a previously published paper ^21^ (**Supplemental Table S2**). **Supplemental Figure S1** presents the representative results of the estimated cell-type maps.

**Figure 1.**
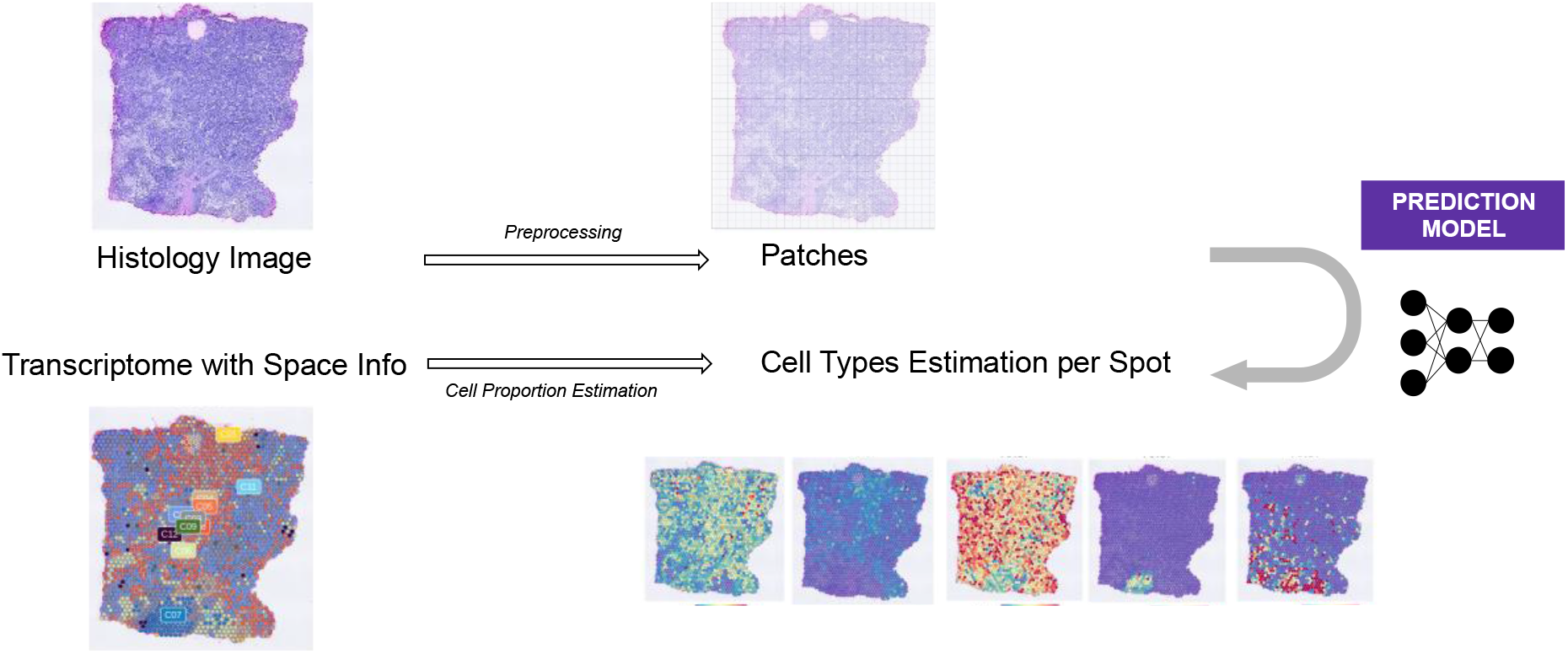
Overview of the deep learning (DL)-based model. We generated spatial transcriptomic data from 22 samples of 10 patients with lung adenocarcinoma (LUAD) containing thousands of spots with transcriptomic data and their spatially matched hematoxylin and eosin (H&E)-stained histologic image. We collected patches of 128 × 128 pixels centered on the spatial transcriptomic spots. The proportion of the five cell types (B cell, NK/T cell, myeloid cell, fibroblast, and epithelial cell) were estimated through the spatial and single-cell RNA-seq data using the cell type inference with domain adaptation (CellDART). A convolutional neural network was trained to predict the enrichment score of the five cell types in each spot.

The correlation coefficient (Pearson’s correlation) between the cell type enrichment scores predicted by the DL model and those of the outputs from the CellDART were calculated. The DL-based scores significantly positively correlated with those of the CellDART across all five cell types as follows: B cell (R = 0.57, *p* < 1 × 10^−5^), T/NK cells (R = 0.29, *p* < 1 × 10^−5^), myeloid cells (R = 0.56, *p* < 1 × 10^−5^), fibroblasts (R = 0.61, *p* < 1 × 10^− 5^), and epithelial cells (R = 0.44, *p* < 1 × 10^−5^) (**Figure 2A**). We also performed validation using independent spatial transcriptomic data (**Figure 2B, C**). The DL-based predicted scores were significantly positively correlated with the scores of the CellDART output across the five cell types (B cell: R = 0.67 and *p <* 1 × 10^−5^; T/NK cell: R = 0.16, *p <* 1 × 10^−5^; myeloid cells: R = 0.18, *p <* 1 × 10^−5^; fibroblasts: R = 0.24, *p <* 1 × 10^−5^; and epithelial cells: R = 0.49, *p <* 1 × 10^−5^).

**Figure 2.**
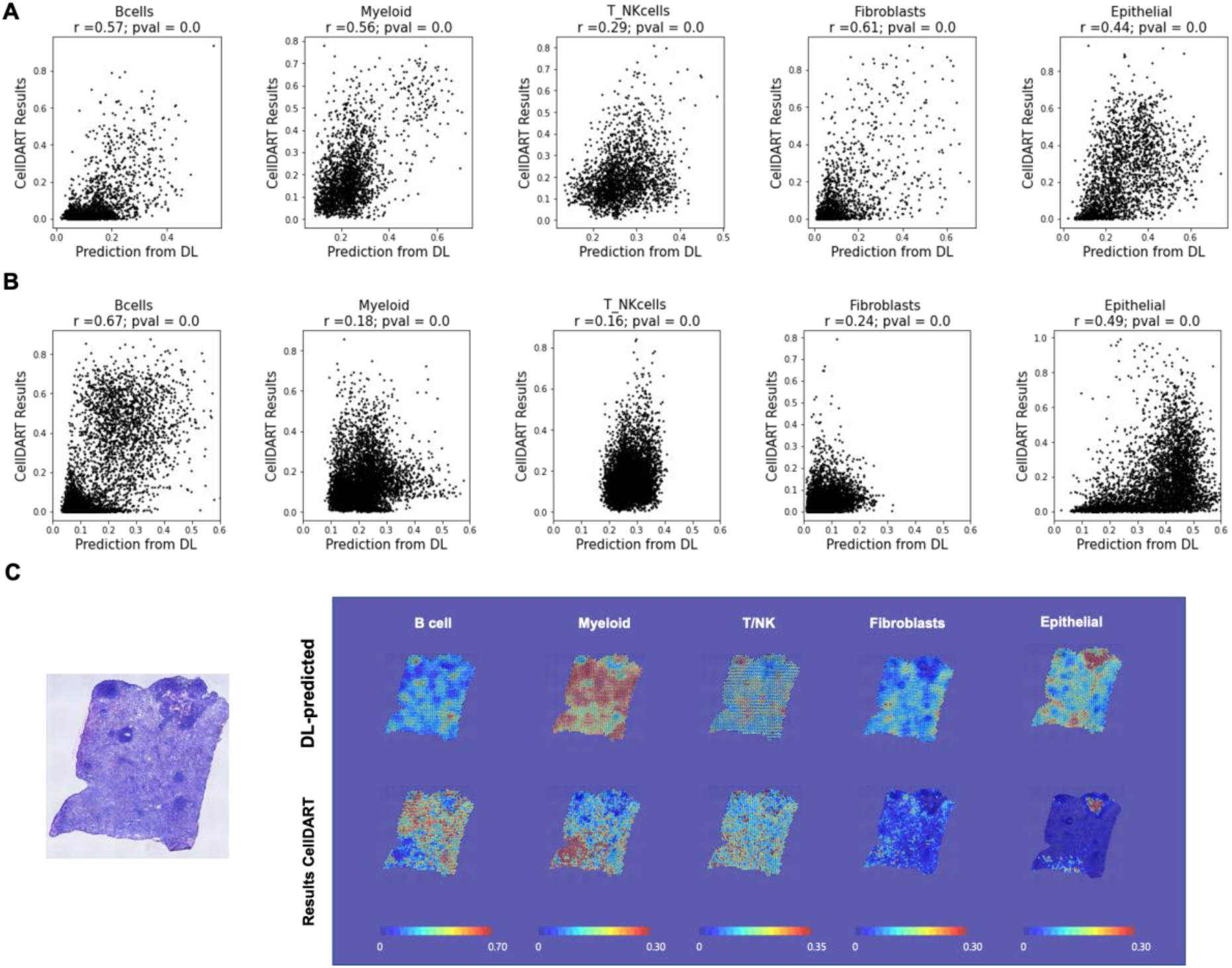
Internal validation of the DL-based model. (A) Patch-wise internal validation using 5% of randomly-selected patches from all patches for model training. Correlations of the five cell types estimation score between the DL model and CellDART on each image patch. (B) Correlations of five cell types estimation score between DL model and CellDART on each image patches from independent spatial transcriptome data. (C) Spatial mapping of the cell type estimation and predicted scores by the CellDART and DL model, respectively, for the five cell types. The outputs were obtained from the one of independent spatial transcriptome samples.

### DL-based and bulk RNA sequencing-based cell types enrichment scores

We compared the enrichment scores of DL-based cell types with those of bulk RNA-sequencing-based cell types to validate the DL-based model further. The H&E and bulk RNA sequencing data of LUAD from TCGA were used. Notably, the regions of tumor samples for bulk RNA sequencing were not directly matched with the regions of H&E images, which is an intrinsic limitation of the TCGA data. We estimated the immune, microenvironment, and enrichment scores of the B cells, macrophages, and cytotoxic lymphocytes from the bulk RNA-sequencing data of TCGA-LUAD samples using the xCell analysis tool ^24^.

We applied the DL model to H&E images of LUAD data to further validate the robustness of the model. To generate heatmaps for the cell type enrichment scores on the H&E slides, the trained DL models were applied to the whole-slide images with a sliding window to estimate the scores for the center of each window (**Figure 3A**). The DL-based scores of 5 cell types of H&E images of TCGA data were calculated (an example is shown in **Figure 3B**). Correlation analyses were performed between the DL-based and bulk RNA-sequencing-based cell-type enrichment scores. **Figure 3C** shows the correlations between the DL-based and bulk RNA-sequencing-based cell type enrichment scores. The DL-based 5-cell type scores were mean values of each tissue. The B-cell estimated using the DL model and RNA-sequencing respectively were significantly positively correlated (R = 0.12, *p =* 0.0076; **Figure 3C**). Myeloid scores of DL-based model also showed a positive correlation with macrophages enrichment scores estimated by RNA-seq (R = 0.15, *p* = 0.001; **Figure 3C**).

**Figure 3.**
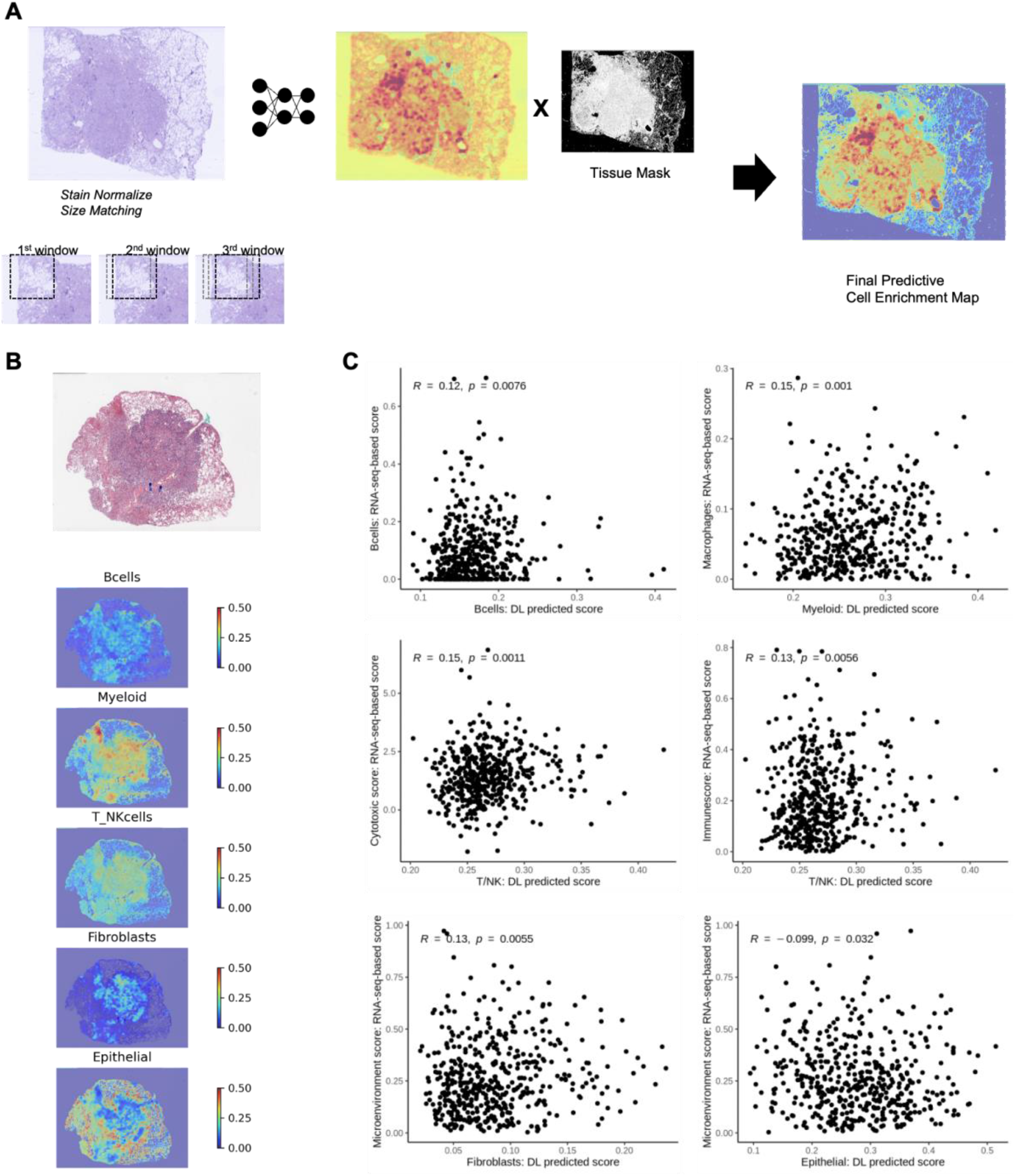
Results from the DL model prediction for the cancer genome atlas (TCGA) - LUAD data. (A) The schematic diagram for predicting five cell types scores from the independent samples with the DL model. The output of the DL model for H&E images was estimated according to the sliding windows, which were reconstructed into a heatmap overlaid with the original H&E image. (B) An example of estimated cell type enrichment score maps using the DL model from a H&E image of TCGA. (C) Scatter plots and correlation coefficients between cell enrichment scores respectively estimated from the bulk RNA-sequencing data and the DL-model were depicted. B cells, myeloid cells, and T/NK cells estimated by the DL model were correlated with B cells, macrophages, cytotoxic score, and immunescore estimated by the bulk RNA-seq data. Microenvironment cell scores were estimated using RNA-sequencing represented other cells excluding cancer cells in the tumor. They were negatively correlated with epithelial scores and positive correlated with fibroblasts estimated by the DL model.

The T/NK cell types estimated by DL-based model were significantly positively correlated with cytotoxic scores (R = 0.15, *p* = 0.001) and immunescore (R = 0.13, *p* = 0.0056) calculated by RNA-sequencing data. Microenvironment scores, which is related to noncancer cells, were positively correlated with DL-based fibroblasts score (R= 0.13, *p* = 0.0055) and negatively correlated with epithelial cells score (R = -0.099, *p* = 0.03; **Figure 3C**).

### External validation on independent LUAD dataset with IHC assay

Accordingly, a heatmap for each cell type was generated from the whole-slide image. Consequently, using the consecutive unstained slides of selected tumor sections, six different antibodies for IHC staining were used as follows: CD3, CD20, CD56, CD68, cytokeratin (CK), and smooth muscle actin (SMA) for the T, B, NK, myeloid, and epithelial cells, and fibroblasts, respectively. **Figure 4** and **Supplemental Figure S2** present the representative samples for this validation process.

**Figure 4.**
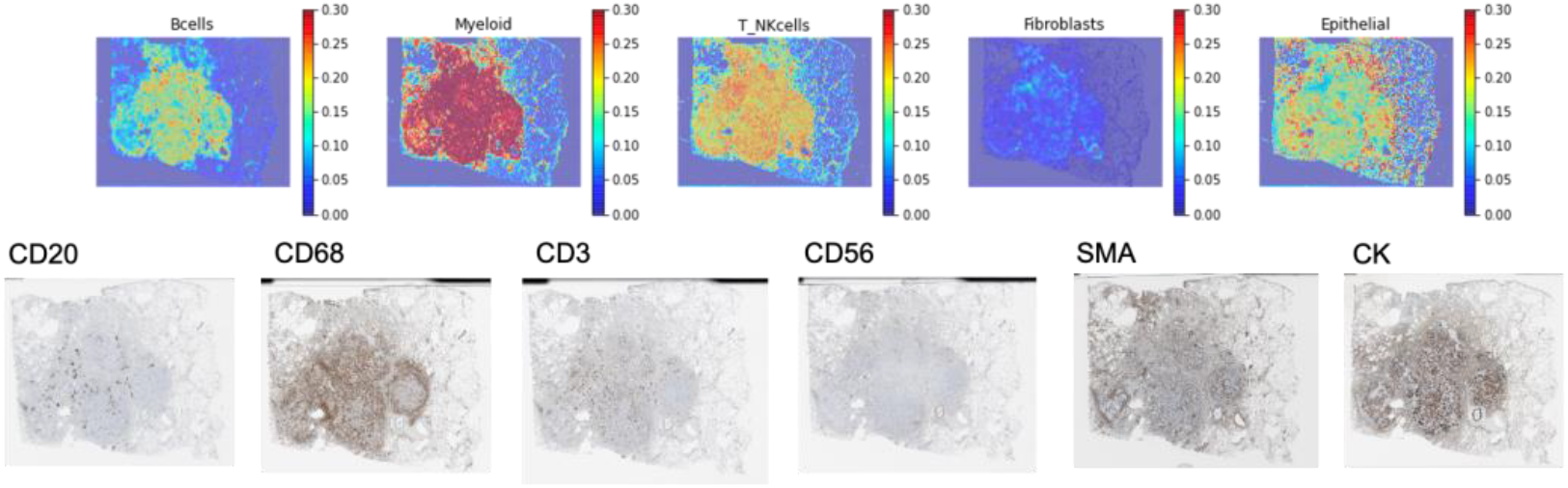
A representative case of DL-based cell type maps and immunohistochemistry (IHC). Visualization of H&E-stained tissue image and predicted cell enrichment score for the five cell types by DL model. IHC staining for CD56, CD20, CD68, CD3, smooth muscle actin (SMA), and cytokeratin (CK) for the same sample were provided.

We performed pixel-wise correlations after spatial registration to compare the results of IHC and DL-based predicted cell-type enrichment score maps. Since H&E and IHC slides were obtained from the adjacent tissue sections, spatial registration was performed using symmetric diffeomorphic image registration ^26^. The DL-predicted cell type enrichment score and deconvoluted image matrix for the DAB channel were applied to Gaussian smoothing, followed by a pixel-wise Spearman’s correlation analysis (**Figure 5A**). As a result, we observed significantly positive correlations between the deconvoluted IHC and DL-based cell type enrichment score as follows: CD20-B (rho = 0.58 ± 0.26; range: 0.09–0.85), CD3-T/NK (rho = 0.66 ± 0.29; range: -0.09–0.91), CD56-T/NK (rho = 0.74 ± 0.15; range: 0.43–0.89), CD68-myeloid (rho = 0.58 ± 0.31; -0.17–0.88), and CK-epithelial (rho = 0.46 ± 0.24; range: 0.12– 0.79) cells, and SMA-fibroblasts (rho = 0.60 ± 0.17; range: 0.34–0.89) (**Figure 5B**). Most of the samples showed strong correlation coefficients between the DL model and IHC assays of the immune cells.

**Figure 5.**
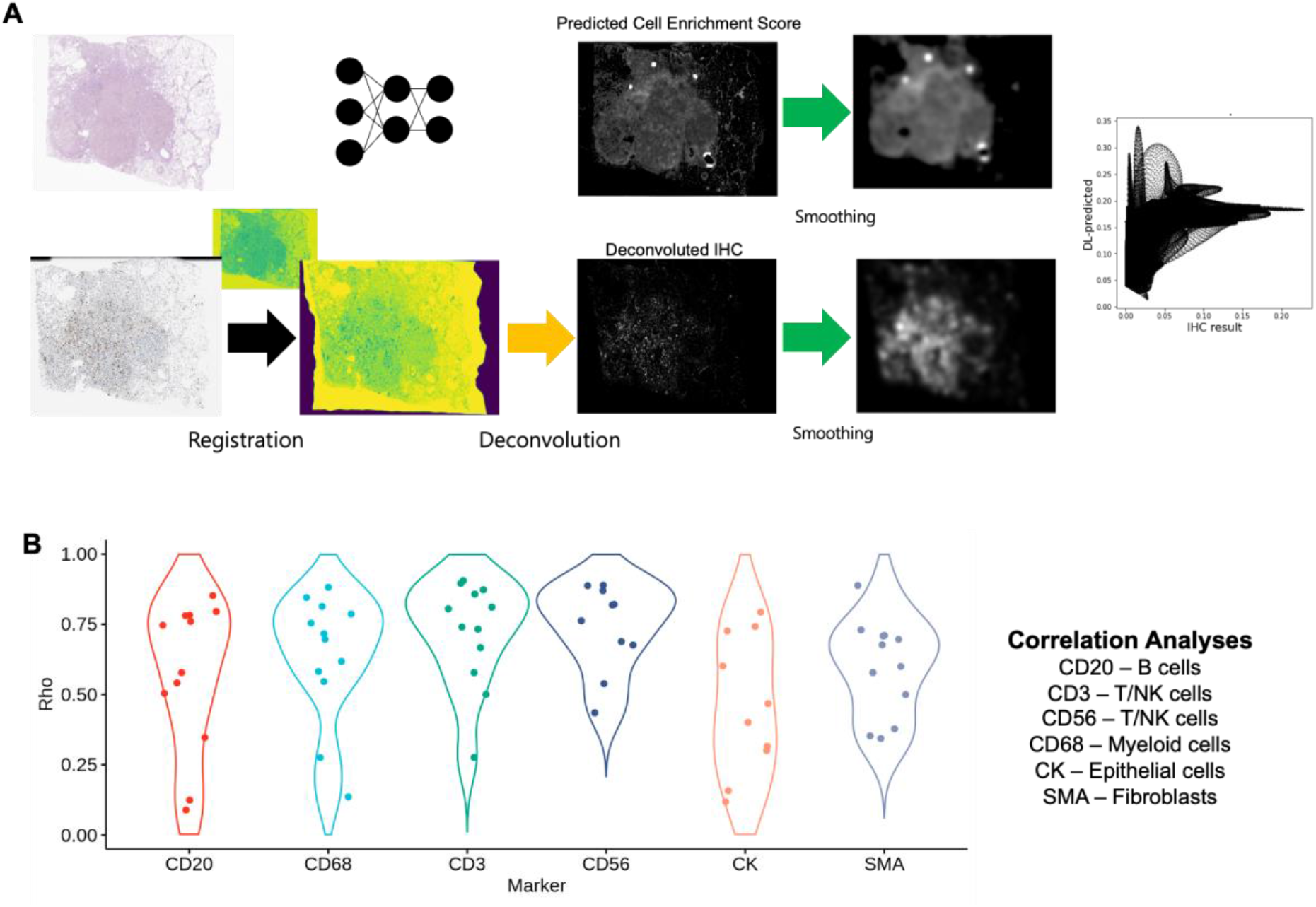
Image-based correlation analysis for external validation. (A) Image registration and deconvolution for IHC performed for the pixel-wise correlation for the IHC map and DL-based output as a heatmap. A 2-dimensional Gaussian smoothing was applied to the deconvoluted IHC images and DL-based output heatmaps to reduce the noise effects linked with fine-level misregistration. The pixel-wise correlation was performed using Spearman’s correlation. (B) Results of correlation analysis between the DL model and IHC staining. (CD20 vs. B cells, CD3 vs. T/NK cells, CD56 vs. T/NK cells, CD68 vs. myeloid cells, CK vs. epithelial cells, and SMA vs. fibroblasts).

## DISCUSSION

Profiling the heterogeneity of TME has been recognized as an essential factor for understanding cancer evolution and developing novel biomarkers and therapeutic agents ^27, 28^. Therefore, RNA-sequencing data as well as histopathologic images have been used to analyze cellular heterogeneity in TME ^29-31^. In particular, one of the major factors in predicting response to ICI is whether immune cells, particularly the T-cells, are enriched in the TME ^32, 33^. However, additional information is required above simple enrichment score of cell types in the TME. Tumor immune properties related to treatment response and prognosis are associated with the spatial contexts of cell types, such as ‘immune-exclusive’ and ‘immune-infiltrative’ types, which is beyond solely quantitative immune cell enrichment ^12^.

This spatial distribution and contexture of multiple cells can be analyzed using histopathologic images. However, the quantitative analysis of the various immune cell types on images remains difficult for several reasons:1) limitations in the labor-intensive labeling process, 2) interpersonal variability, and 3) only a few cell types that can be visually identified. Notably, integrating RNA-sequencing data with high-resolution and spatially co-registered information is a major opportunity for spatial transcriptomics technology.

Consequently, investigators could interpret the complex spatial characteristics of the TME and develop computational models by combining transcriptome data, spatial location, and visual features of cell morphology. Using this integrative approach, we illustrated that a readily available H&E histopathological image could predict the spatial distribution of major cell types without manual labeling and employing intrinsic data features. We found that the cell proportion scores predicted by the DL model from the histopathological images had a positive correlation with cell estimation scores calculated using the bulk RNA-sequencing data from the TCGA data. We also used IHC assays to validate the model, which was accurate in the independent surgical cohorts.

This DL model could be used for various purposes in clinical settings, particularly to examine immunotherapy response and prognosis while considering the immune contexts using H&E images alone. Spatial patterns of specific immune and stromal cells are linked with the responses to ICIs and clinical outcomes ^34^. Moreover, a recent study illustrated that the spatial phenotype of tumor-infiltrating lymphocytes within the TME using an artificial intelligence-powered model showed a significant association with immunotherapy response in non-small cell lung cancer ^18^. Since our model could determine B-cell enrichment maps in the TME, the location and localized patterns of these cell types can aid us in identifying tertiary lymphoid structures ^35^. Notably, the presence of tertiary lymphoid structures in solid tumors is associated with immunotherapy response and progression-free survival ^36-38^.

Therefore, we believe the spatial phenotype using our approach could be used as a method to develop a novel biomarker for prognostic stratification and treatment response in various solid tumors. In addition, as various transcriptomic analysis has been used to find new molecular markers associated with prognosis and response, our framework using a DL model could be trained to predict these markers or molecular signatures on spatial transcriptomic data. Then, this DL model can predict key molecular markers only using the histologic images.

The strategy that uses major cell types or molecular pathways derived from spatial transcriptomics as labels for developing DL models could be used to develop flexible biomarkers for companion diagnostics with numerous potential therapeutics for the TME. In precision medicine approaches, various biomarkers are extensively adopted in clinical trials to apply new therapeutics to select patients who are expected to benefit from the therapeutics ^39^. Recently, novel therapeutic targets for immune and stromal cells within the TME, as well as T-cells, have been developed and assessed in phase 1 or 2 clinical trials ^40, 41^. Similar to our suggested strategy, cell type maps were constructed based on scRNA-seq data, and the model could show the specific cell types or molecular markers within the TME with their spatial contextures ^42^. For example, novel therapeutics that target tumor-associated macrophages or fibroblasts can be analyzed using a potential biomarker derived from the proposed DL model.

Conclusively, our study showed that integrating cell-type maps derived from spatial transcriptomics and histopathologic images using DL facilitated the prediction of TME characteristics in LUAD. This strategy will enable various types of image-based biomarkers to demonstrate spatial phenotypes that could be used in clinical settings, such as prognosis stratification and predicting response to therapeutics targeting tumor microenvironment.

## Ethics statements

### Ethics approval

This study was approved by the institutional review board of our institute, and informed consent was obtained from all patients. All procedures performed in this study involving human participants followed the ethical standards of the institutional and/or national research committee and the 1964 Helsinki declaration and its later amendments or comparable ethical standards.

### Patient consent for publication

*Not applicable*.

