## Supplementary Figure 1 for "Mapping cell types in the tumor microenvironment from tissue images via deep learning trained by spatial transcriptomics of lung adenocarcinoma"

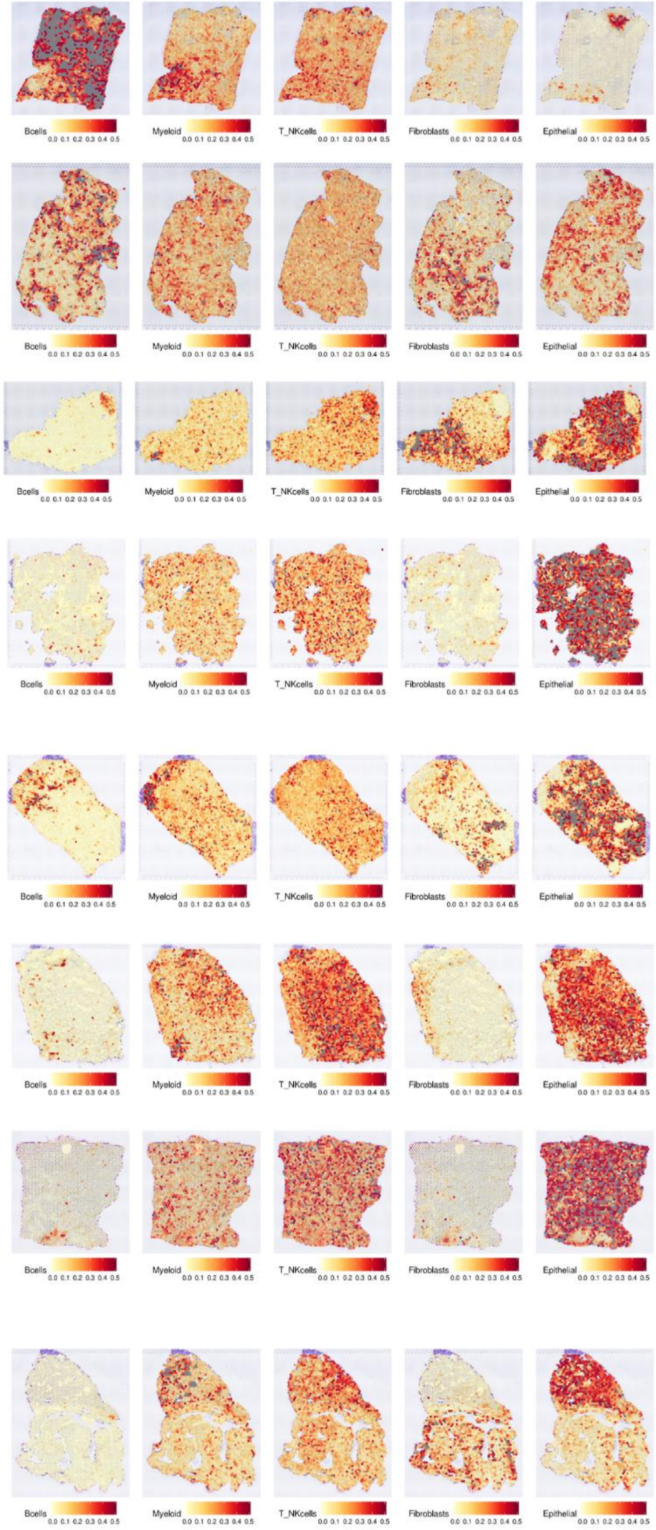


**Supplementary Figure 1.** Results of CellDART to estimate five cell types defined by single cell-RNA-seq to map spatial transcriptomics data of lung adenocarcinoma
