## Supplementary figures and images for "Mapping cell types in the tumor microenvironment from tissue images via deep learning trained by spatial transcriptomics of lung adenocarcinoma"

### Supplementary Figure 2

**Figure S2. Examples of DL-based cell type mapping and immunohistochemistry using key markers**

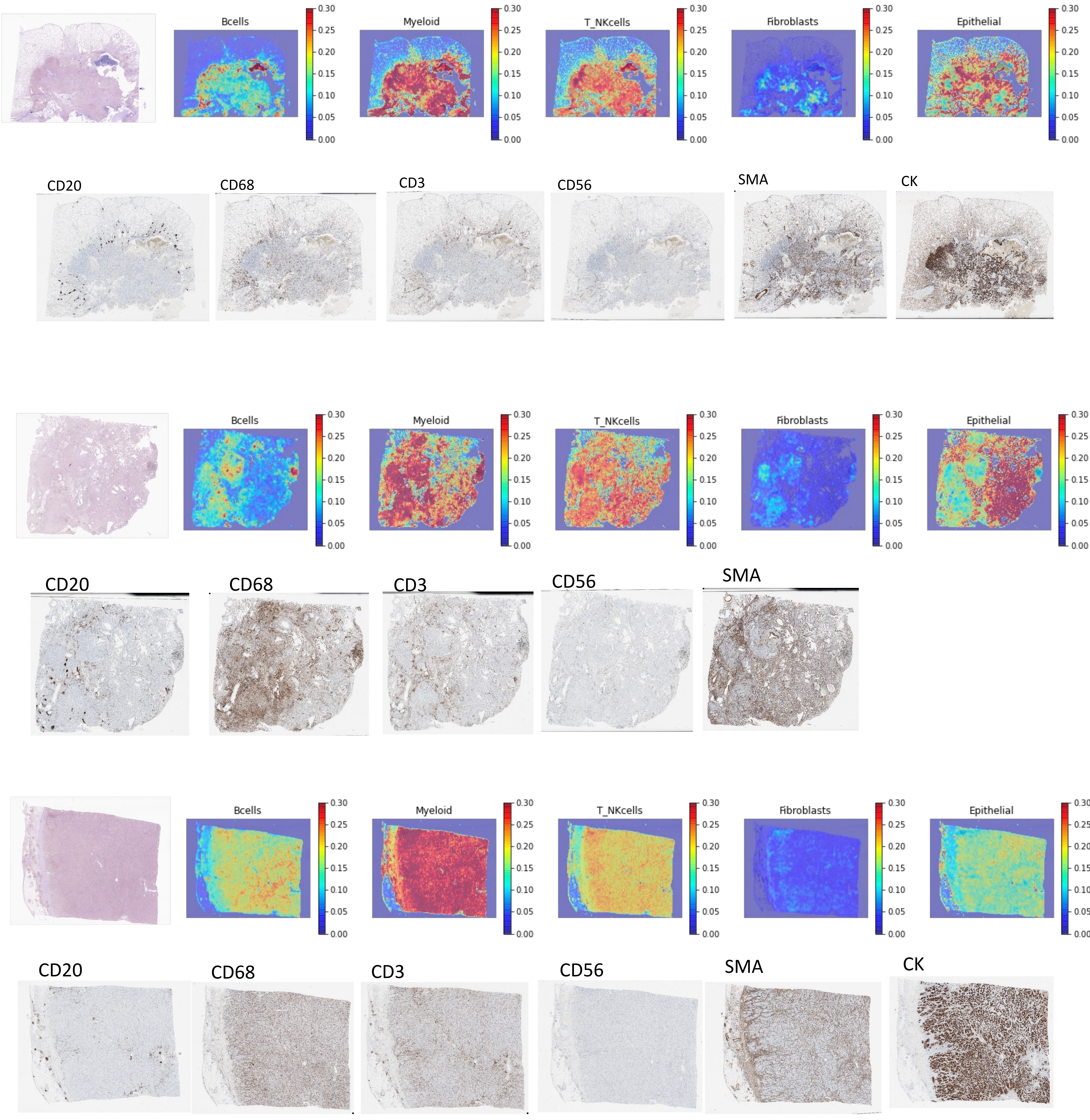
